# Macrophage-specific NF-κB activation dynamics can segregate inflammatory bowel disease patients

**DOI:** 10.1101/535096

**Authors:** Stamatia Papoutsopoulou, Michael D. Burkitt, François Bergey, Hazel England, Rachael Hough, Lorraine Schmidt, David G Spiller, Michael HR White, Pawel Paszek, Dean A. Jackson, Vitor A.P. Martins Dos Santos, Gernot Sellge, D. Mark Pritchard, Barry J. Campbell, Werner Müller, Chris S. Probert

## Abstract

The heterogeneous nature of inflammatory bowel disease (IBD) presents challenges, particularly when choosing therapy. Activation of the NF-κB transcription factor is a highly-regulated, dynamic event in IBD pathogenesis. We expressed the human NF-κB/p65 subunit in blood-derived macrophages, using lentivirus. Confocal imaging of p65 activation revealed that a higher proportion of macrophages from Crohn’s patients responded to lipid-A compared to controls. In contrast, cells from ulcerative colitis (UC) patients exhibited a shorter duration of p65 nuclear localisation compared to healthy controls and Crohn’s donors. Using a similar lentivirus approach, NF-κB-regulated luciferase was expressed in patient macrophages, isolated from frozen peripheral blood mononuclear cell samples. Following activation, samples could be segregated into three clusters based on the NF-κB-regulated luciferase response. The majority of UC samples appeared in hypo-responsive cluster 1, with Crohn’s patients representing the majority of hyper-responsive cluster 3. A positive correlation was seen between NF-κB-induced luciferase activity and cytokine levels released to medium from stimulated macrophages, but not in serum or biopsy. Analysis of macrophage cytokine responses and patient metadata revealed a strong correlation between Crohn’s patients who smoked and hyper-activation of p65. These *in vitro* dynamic assays of NF-κB activation in blood-derived macrophages segregate IBD patients into groups with different phenotypes and therefore may help determine response to therapy.

**Significance statement:** This manuscript describes two dynamic assays of NF-κB activation in blood-derived macrophages that can segregate IBD patients into groups with different phenotypes. For the first time we introduce the use of dynamic measurements of a transcription factor activation as a method to stratify patients and we are confident that our approach will lead in future to early patient stratification and prediction of treatment outcome.

## Introduction

Inflammatory bowel disease (IBD) is characterised by an imbalanced immune response, leading to a pro-inflammatory phenotype with elevated tissue concentrations of various cytokines including tumour necrosis factor (TNF), interleukin-6 (IL-6) and interferon-γ (IFNγ) (1,2). One of the key mechanisms involved in generating this inflammatory environment within the intestinal mucosa is the activation of the transcription factor nuclear factor kappa-light-chain-enhancer of activated B cells (NF-κB) (3). The NF-κB family of transcription factors contains five subunits, p65 or RelA, p50, c-Rel, p52 and RelB, that may function as homo- or hetero-dimers. These dimers are retained in an inactive state in the cytoplasm by binding to a member of the family of inhibitory κB proteins (IκBs). Upon stimulation, IκBs are degraded, and the NF-κB active dimers translocate into the nucleus where they regulate transcription. The nuclear translocation of NF-κB proteins is a highly regulated, dynamic event, characterised not only by the transport of NF-κB dimers into the nucleus following stimulation, but also by shuttling between the nucleus and cytoplasm of the cell, with context-specific oscillatory frequency; such as has been observed for transcriptionally active p65 subunit dimers (4). To date, these heterogeneous dynamics have been demonstrated using mouse (5-7) and human cell-lines (8), and primary cells obtained from transgenic mice expressing fluorescent fusion proteins (9,10).

NF-κB activation and dysregulated cytokine production has previously been reported in various cell types in IBD patients (11,12). Macrophages and epithelial cells isolated from inflamed intestinal biopsies showed augmented levels of NF-κB (13). Lamina propria fibroblasts have been reported to be involved in cytokine production due to highly activated p65 (14), and epithelial NF-κB signalling has been implicated in several murine models of IBD (11,15,16).

Inevitably, the NF-κB pathway has therefore become an attractive target for therapeutic interventions in IBD, and many of the current drugs that are used to treat IBD either directly or indirectly influence NF-κB signalling (e.g. corticosteroids, anti-TNF monoclonal antibodies and 5-aminosalicylates). Nevertheless, a significant proportion of patients do not respond to these treatments (17-19). The reasons for treatment failure are not completely understood. In clinical practice, there is an increasing use of therapeutic drug monitoring to characterise treatment failure, but, even in carefully observed cohorts, the development of anti-drug antibodies can only explain a small proportion of secondary losses of response to therapy and does not explain primary failure to respond (20).There is, therefore, a need for more specific laboratory tests to better stratify patients for therapy prior to drug initiation. We hypothesised that patients with different clinical phenotypes would have differences in their NF-κB activation responses. Previous studies looking at NF-κB activation in human samples have largely relied on static measurements of DNA-binding activity using electrophoretic mobility shift assays (EMSA) (21). Whilst EMSAs clearly demonstrate NF-κB DNA binding, they do not demonstrate transcriptional effects of DNA binding, and are thus unable to illustrate the dynamics and heterogeneity of NF-κB activation. Hence, in this study, we describe a novel screening protocol based on the dynamic detection of endogenous NF-κB activation in peripheral blood mononuclear cell-derived macrophages. We report the characteristics of this assay, including its reproducibility and its ability to segregate individuals into different groups. Changes in NF-κB dynamics correlated with differences in cytokine response and we report how this reflects the clinical phenotype of individuals.

## Results

### Heterogeneity of NF-κB activation reveals differences between IBD patients and healthy controls using peripheral blood monocyte-derived macrophages isolated from a pilot cohort

Macrophages play a pivotal role in the pathogenesis of IBD. In this context, they have been shown to acquire a pro-inflammatory phenotype (M1) and secrete cytokines. This process is critically regulated by the NF-κB family of transcription factors (22). NF-κB dimers, such as p50/p65, are activated in pro-inflammatory macrophages, regulating the expression of cytokines and chemokines. In this study, we used a lentivirus system to express human p65-AmCyan in primary PBMDM to determine whether there was an alteration in p65 nuclear translocation dynamics in cells from patients compared to control volunteers. We recruited a pilot cohort of 14 patients from the Royal Liverpool and Broadgreen University Hospitals NHS Trust, who donated fresh peripheral blood from which PBMDMs were generated for confocal imaging of NF-κB activation (Figure 1A). Dynamic p65 nuclear translocation within lentivirus-transduced cells stimulated with Lipid A, a main cytotoxic component of LPS of Gram-negative bacteria (23), could be clearly visualized by time-lapse confocal microscopy (Figure 1B). Overall, we screened 118 cells from healthy controls (n=6 donors), 59 cells from CD patients (n=5 donors) and 30 cells from UC patients (n=3 donors). In resting cells, p65-AmCyan was localized almost exclusively in the cytoplasm within all cells but exhibited rapid nuclear translocation within minutes of stimulation. Notably, only a single p65 translocation (rather than repeated shuttling) was observed (within the 3h period). While there was considerable variation between single cell responses, the average nuclear p65-AmCyan trajectory, as well as the amplitude and timing were similar in samples taken from IBD patients and controls (Figures 1C, 1D and 1E). Previously, NF-κB dynamics have been shown to exhibit “all-or-nothing” activation, where only a fraction of cells in the population exhibited translocation following low dose stimulation (6,24), temporal cytokine pulses (8) or heat stress (25). We, therefore, quantified the fraction of cells that exhibited p65-AmCyan translocation in samples taken from patients and healthy controls (Figure 1C). This suggested that the dynamics of the p65 response, at least in the screened samples, did not differ between groups. Altogether, out of >200 single cells that were assayed across all conditions approximately 40% of cells responded to Lipid A stimulation across individuals in the healthy control group, although a significantly higher proportion of cells (61%) were responsive in the CD patient group; p<0.001; two-sided Fisher’s exact test by summation (Figure 1F; Supplementary Information: Table S1). Inspecting the characteristics of p65-amCyan translocation in responding cells, we found that this did not differ between groups in terms of the average trajectory, timing and amplitude (Figures 1G and 1H, respectively). Interestingly though, the peak width, which reflects the duration of NF-κB nuclear localization, was observed to be significantly lower in responding cells from UC patients compared to either controls or CD patients (Figure 1I; p<0.001; Kruskal Wallis test).

**Figure 1:**
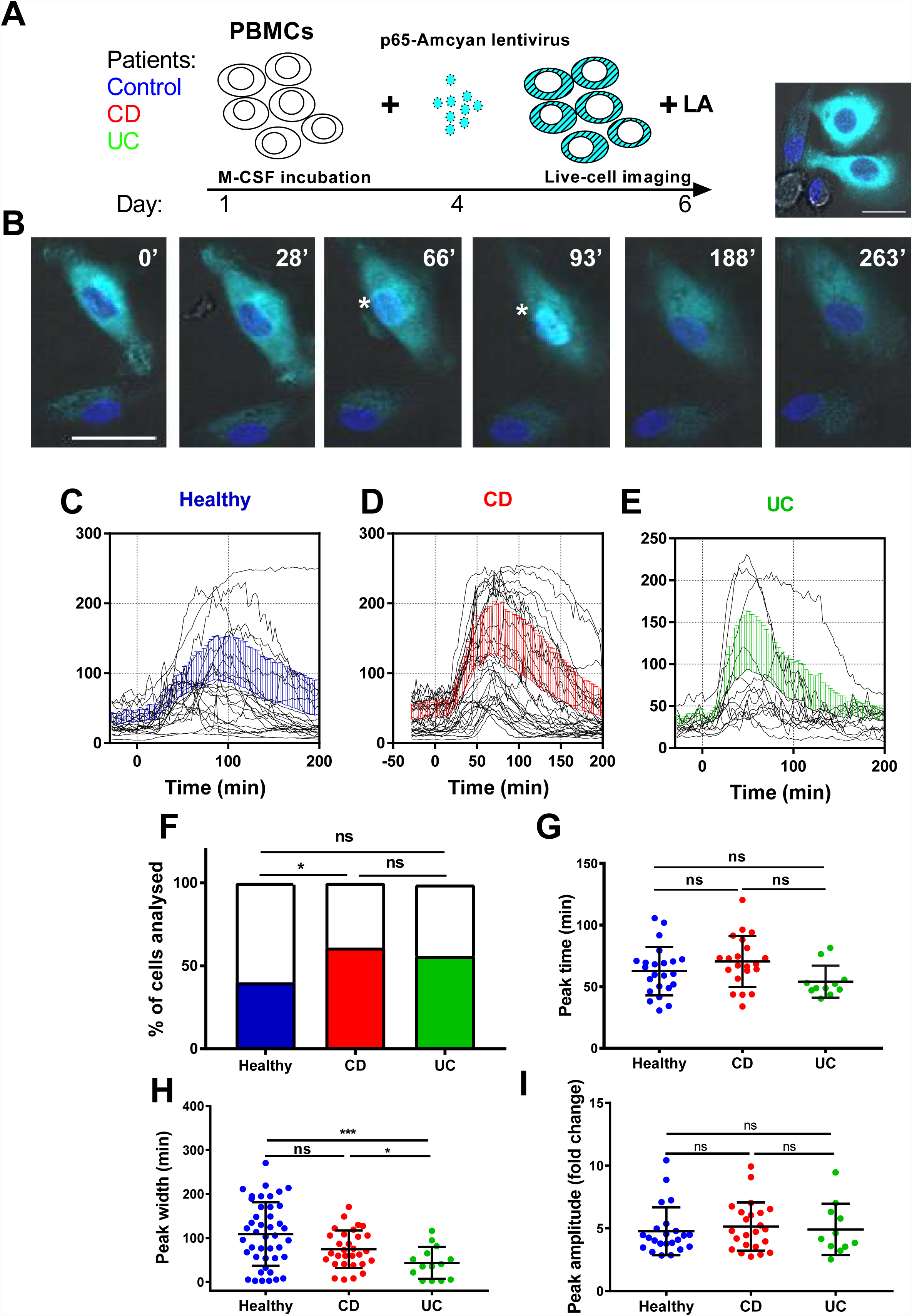
Time-lapse Confocal imaging of NF-κB nuclear translocation in Lipid A-stimulated human PBMDMs. (A) PBMC-derived macrophages from human subjects were transduced with a fluorescent lentiviral p65AmCyan construct and were used to image NF-κB p65 subunit cellular dynamics in real-time. (B) CellTracker was used to generate profiles of nuclear translocation of transduced cells post stimulation with 200ng/mL Lipid A, over 3h: Asterisks (*) indicate nuclear localization of p65 at 66 and 93 min (‘). (C-E) Graphs of individual nuclear translocation profiles of responding cells in the healthy controls, Crohn’s disease (CD) and ulcerative colitis (UC) groups; mean ± SD for all responsive cells shown in blue, red and green colour lines for each group, respectively. (F) The proportion of responding cells was calculated by pooling all tracked cells from each disease group; *p<0.05; two-sided Fisher’s exact test (by summation). Peak identification and analysis revealed in each population the peak time (G), peak amplitude (H) and peak width (I): *p<0.05, ***p<0.001; statistical differences assessed by Kruskal Wallis test with pairwise comparison of all three groups.

### Patient demographics of the main cohort study to examine NF-κB activity

As part of the SysMedIBD project (https://www.sysmedibd.eu/) a novel cohort of patients was recruited from two centres in Western Europe. In this study, we have investigated NF-κB signalling in 65 subjects, including healthy donors (Control) and patients with CD or UC (Table 1). UC patients were found to be significantly older than those with CD (p=0.03; Mann-Whitney-Wilcoxon test) and Control subjects (p=0.01) and to have a higher BMI than patients with CD (p=0.04). CD patients also had a higher serum CRP concentration than those with UC (p=0.03). No differences were detected in terms of smoking status, gender, or use of immunomodulators (thiopurines, methotrexate) or biologics (infliximab, adalimumab, vedolizumab or ustekinimab).

**Table 1:** Participants for luciferase-based screening studies were recruited from outpatient clinics at The Royal Liverpool and Broadgreen University Hospitals NHS Trust, UK and University Hospital Aachen, Germany.

| Disease status | Crohn's disease | Ulcerative colitis | Control | p-value CD vs UC | p-value CD vs C | p-value UC vs C |
| --- | --- | --- | --- | --- | --- | --- |
| Number of patients | 26 | 14 | 25 |  |  |  |
| Gender, (% male) | 50% (13/26) | 64.3% (9/14) | 40% (10/25) | 0.594 | 0.663 | 0.262 |
| Age Mean (SD) | 38.3 (14.1) | 48.5 (14.1) | 35.3 (12.3) | 0.033 | 0.534 | 0.010 |
| BMI Mean (SD) | 24.2 (3.2) | 28 (5.3) | 27.6 (6.7) | 0.044 | 0.229 | 0.966 |
| Smoking (%)<sup>1</sup> |  |  |  |  |  |  |
| Current | 30.8% (4/26) | 7.7% (1/13) | 12.5% (3/24) | 0.227 | 0.224 | 1 |
| Previous | 15.4% (8/26) | 15.3% (2/13) | 20.8% (5/24) | 1 | 0.895 | 1 |
| Never | 53.8% (14/26) | 77% (10/13) | 66.7% (16/24) | 0.295 | 0.525 | 0.783 |
| CRP Mean (SD) | 7.3 (14) | 1.7 (3.6) | 2.1 (3.5) | 0.031 | 0.799 | 0.088 |
| Immunomodulator (thiopurines, methotrexate) % | 23.1% (6/26) | 14.3% (2/14) |  | 0.803 |  |  |
| Biologic drug prescription (infliximab, adalimumab, vedolizumab, ustekinumab) % | 30.8% (8/26) | 28.6% (4/14) |  | 1 |  |  |
| Disease behaviour (%)<sup>2</sup> |  |  |  |  |  |  |
| Penetrating disease (B3) | 24% (6/25) |  |  |  |  |  |
| Stricturing disease (B2) | 24% (6/25) |  |  |  |  |  |
| Neither stricturing, nor penetrating disease (B1) | 52% (13/25) |  |  |  |  |  |

1 Missing data for one Ulcerative colitis patient and for one Control
2 Missing data for one Crohn's disease patient

### Measurement of NF-κB activity in PBMDMs is reproducible and demonstrates inter-individual variability

To characterize NF-κB activation in macrophages from our main cohort of patients and donors, isolated monocytes from PBMC samples were differentiated into PBMDMs and subsequently transduced with a lentiviral construct that expresses firefly luciferase under the control of the classical NF-κB promoter. Cultures were activated by addition of LPS, and luciferase activity was monitored over time. To investigate the dynamics of this assay, we performed LPS dose-response studies. These demonstrated a strong luciferase response to LPS at a dose of 200ng/mL LPS. However, marked differences in luciferase response were observed between individuals (Figure 2).

**Figure 2:**
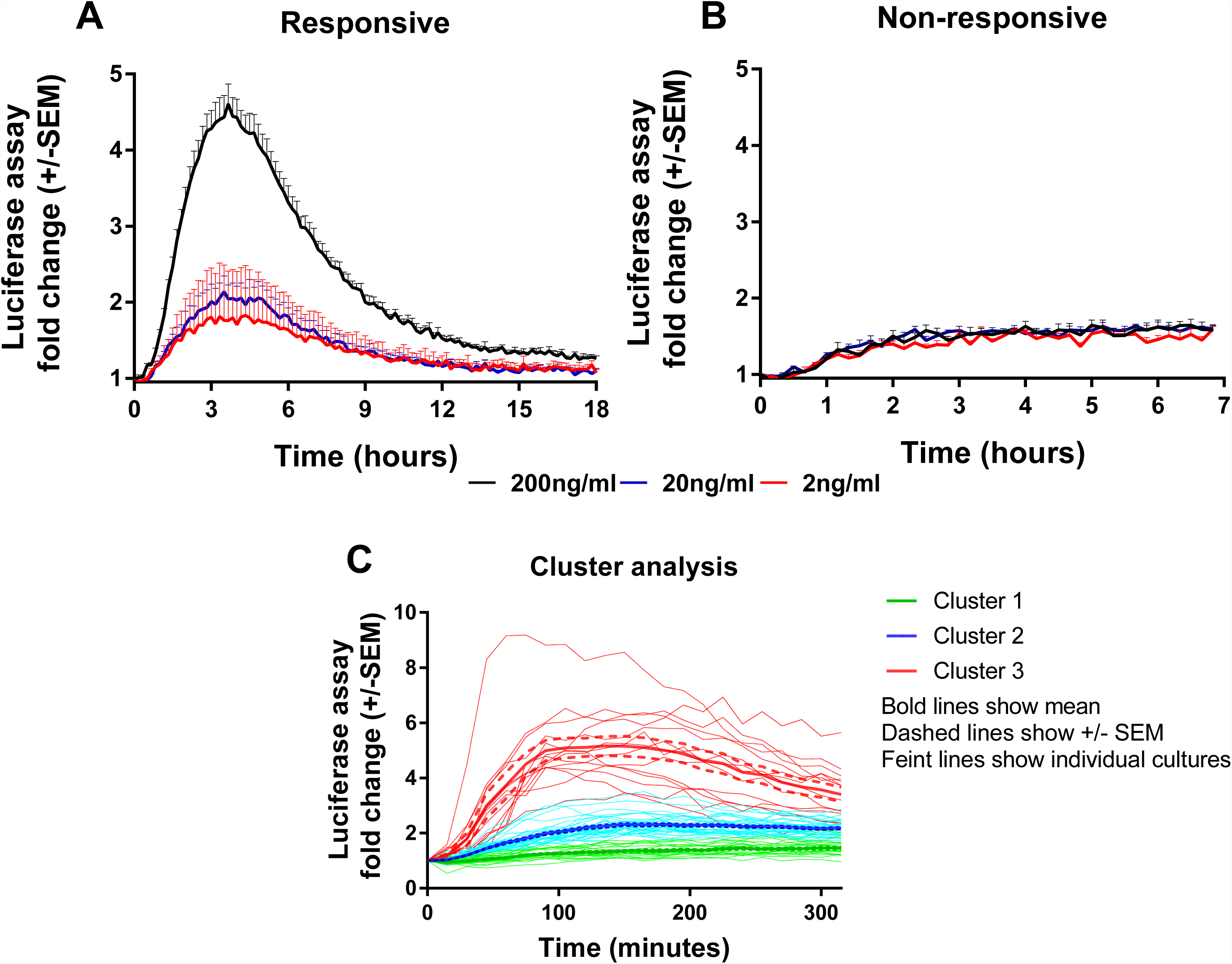
Endogenous NF-κB-regulated luciferase activity in LPS-stimulated PBMDMs. Differentiated PBMDMs (5 vials per patient) were stimulated with increasing doses of LPS; 2ng/mL (red line), 20ng/mL (blue line) and 200ng/mL (black line) and luciferase activity was measured over time. Graphs of a representative patient are shown in Responsive (A) or Non-responsive samples (B). LPS-stimulated PBMDMs were assigned to one of three clusters based on log2 fold change in the luciferase activity profile using K-medoids algorithm (C).

To determine whether this phenotypic variation was attributable to an extra-corporeal effect, we undertook validation studies in which we used multiple vials (five) of PBMCs isolated from the same individual. Cells from each vial were thawed every week and differentiated, transduced and stimulated independently (Figure 2A and 2B). These data demonstrated reproducibility of luciferase activation plots, suggesting that the differences observed between samples from individuals represent biological differences, rather than being attributable to differences in culture technique.

To determine whether the luciferase response was affected by alternative sources of LPS, we used κB-NLSluc-transduced PBMDM samples from 12 randomly selected individuals from the CD cohort and stimulated them with doses of either commercial LPS from *Salmonella* Minnesota, or with LPS extracted from two CD disease mucosa-associated adherent, invasive *E. coli* strains, LF10 and LF82. All sources of LPS that were tested induced a similar dose-dependent luciferase response (Supplementary information: Figure S1A). In addition, we also investigated the impact of various other stimulatory ligands on luciferase activity: Muramyl-dipeptide (MDP; an activator of Toll-like receptor TLR2), flagellin (for TLR5), and IL-1β (for IL-1R), all induced NF-κB activation, but none of these stimuli were found to be as potent as LPS (Supplementary information: Figure S1B).

Having demonstrated the reproducibility of luciferase response to LPS, we screened PBMDMs derived from all patients to determine their luciferase responses. Patients were stratified according to their log_2_ fold change of luciferase activity using K-medoids algorithm to assign each sample to one of three clusters. Cluster 1 contained samples from the least responsive group. Cluster 3 contained samples from the most responsive group, whilst cluster 2 contained the samples that fell between these two extremes (Figure 2C). Samples from both Liverpool and Aachen were represented in each cluster and cells from healthy donors from both locations showed similar behaviors (Supplementary information: Figure S1).

### High luciferase activity reflects a strong pro-inflammatory phenotype

To investigate the biological consequences of the differences in NF-κB activation observed, we measured pro-inflammatory cytokine concentrations produced by LPS-induced PBMDMs from the two extreme clusters, cluster 1 and cluster 3. For this purpose, we identified 8 individuals from Cluster 1, and the same number from Cluster 3, who had also undergone colonoscopy and biopsy of areas of macroscopically normal tissue from both the terminal ileum and sigmoid colon. New cultures of PBMDMs were prepared from each individual and these were stimulated with 200ng/mL LPS for 20h. The culture medium was harvested to measure the concentrations of several pro-inflammatory cytokines. Our initial analysis of this data was to perform a principal component analysis (PCA) and this demonstrated a clear separation between Cluster 1 and Cluster 3 based on stimulated cytokine secretion (Figure 3A). The differences demonstrated by PCA analysis are also reflected in analysis of the concentrations of individual cytokines. PBMDMs from cluster 3 produced substantially higher levels of TNF (p=0.01; Mann-Whitney U test), IL-1β (p<0.001), IL-10 (p<0.001) and IL-8 (p=0.04) compared to Cluster 1 (Figure 3B) on univariate analysis. When corrections were made for multiple testing using the FDR method, the differences observed for TNF, IL-1β and IL-10 remained statistically significant.

**Figure 3:**
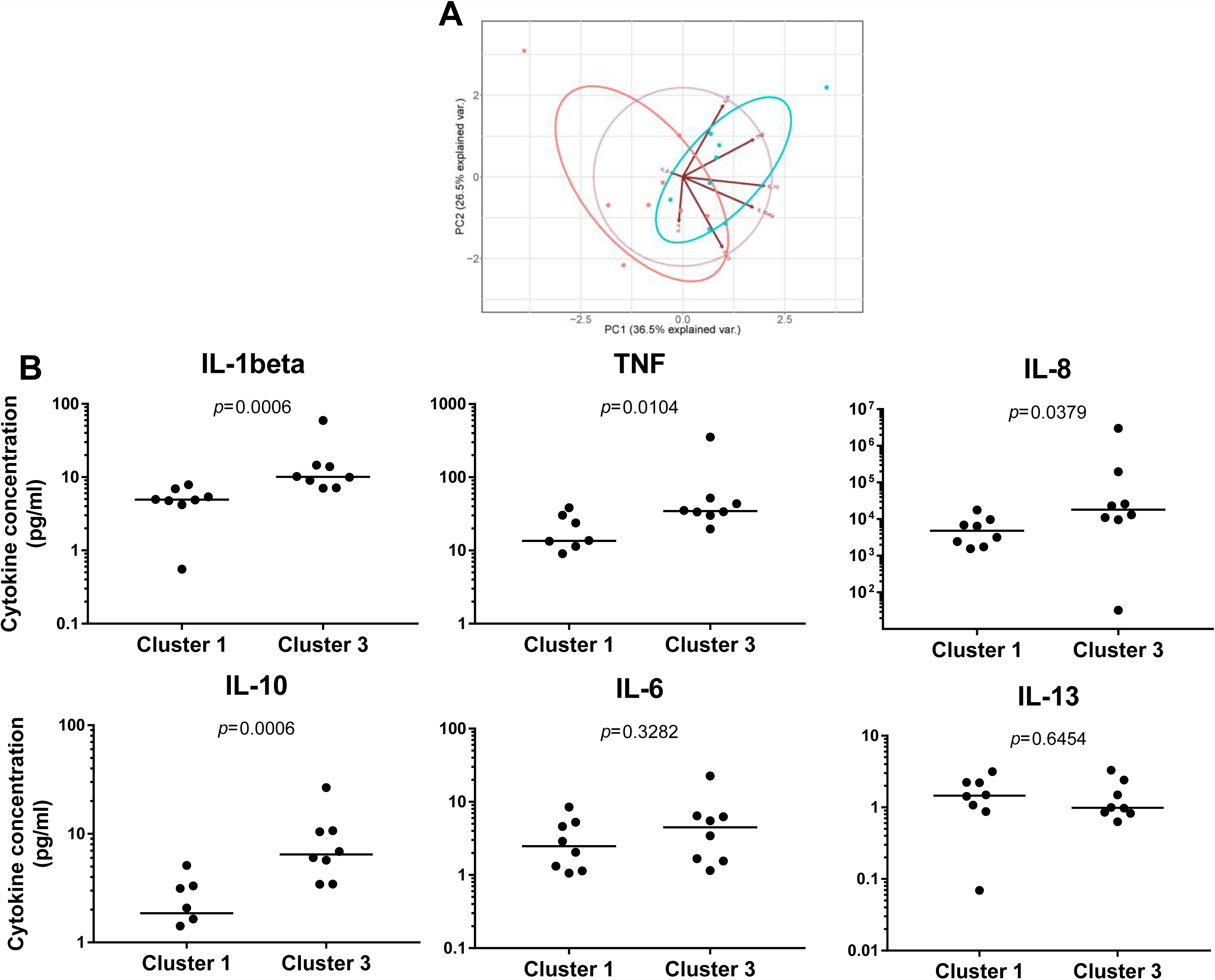
Pro-inflammatory cytokine levels can reflect differences between NF-κB-based clusters. PCA analysis based on concentrations of pro-inflammatory cytokines secreted in medium of LPS-stimulated PBMDMs from cluster 1 and cluster 3 samples (A). Quantification of individual cytokines from this analysis. (B). Statistical comparisons between clusters were made using the Mann-Whitney-U test, individual p-values are reported on each chart.

To determine whether these differences in cytokine production were also reflected in the serum or enteric mucosa of the patients from whom the PBMDMs had been derived, we performed cytokine assays using serum, as well as homogenised tissue lysates from the terminal ileum and sigmoid colon of the same patients (Supplementary Information: Figure S2). In the serum, IL-1β levels were undetectable (dynamic range 0.05-375 pg/mL), but the concentrations of the other cytokine levels that were examined were found to be similar in between the two clusters. Similarly, few differences were observed in cytokine levels in intestinal tissue homogenates between patients from each the two cluster. The only exception was a marginal, but significantly lower abundance of IL-13 observed in the lysates of sigmoid colon tissue obtained from patients in Cluster 3 (Cluster 1; median 9.9 pg/mL (IQR, 7.4-11.4) vs Cluster 3 = 5.3 pg/mL (IQR, 4.4-7.5); p=0.0401, Mann-Whitney U test). However, this difference was not statistically significant when p-values were corrected for multiple comparison by FDR. Further comparison of the levels of each cytokine from LPS-stimulated PBMDMs (dynamic measurement) and intestinal tissue samples (static measurement) also revealed no correlation between dynamic and static measurements (Supplementary information: Figure S3). This suggests that the dynamic measurement of *in-vitro* cytokine production likely reflects the increased activity of endogenous NF-κB and the pro-inflammatory status of the PBMDMs.

### Low NF-κB regulated luciferase activity is characteristic of ulcerative colitis patients

Within the three distinct clusters representing differential luciferase activity, we noticed that Cluster 1, which showed very low luciferase activity, included mainly samples that had been obtained from patients with UC. We therefore analysed luciferase activity data obtained from patient samples taken from both clinical sites (Liverpool and Aachen) and based on disease status (Figures 4A and B). A marked difference in luciferase activity from PBMDMs in response to stimulation with 200ng/mL LPS over 20h was observed only between healthy donors (controls) and UC samples and not with CD, with UC samples showing significantly lower levels of luciferase activity shows; p<0.05, Kruskal-Wallis test (Figure 4B). On the other hand, PBMDMs derived from CD patients were represented in in all three assigned clusters, with a very broad spectrum of luciferase activity being observed upon stimulation with LPS. Further analysis of the metadata revealed that PBMDMs from CD patients who were active smokers showed significantly higher luciferase activity than those from non-smokers (Figure 4C; *p*<0.05, ANOVA). Interestingly, this was not the case for the control sample LPS-stimulated PBMDMs, where there was no observed difference between smokers and non-smokers; *p*=0.4516 (Figure 4D). Most patients with UC are non-smokers hence there were few samples available from UC patients who were also current smokers.

**Figure 4:**
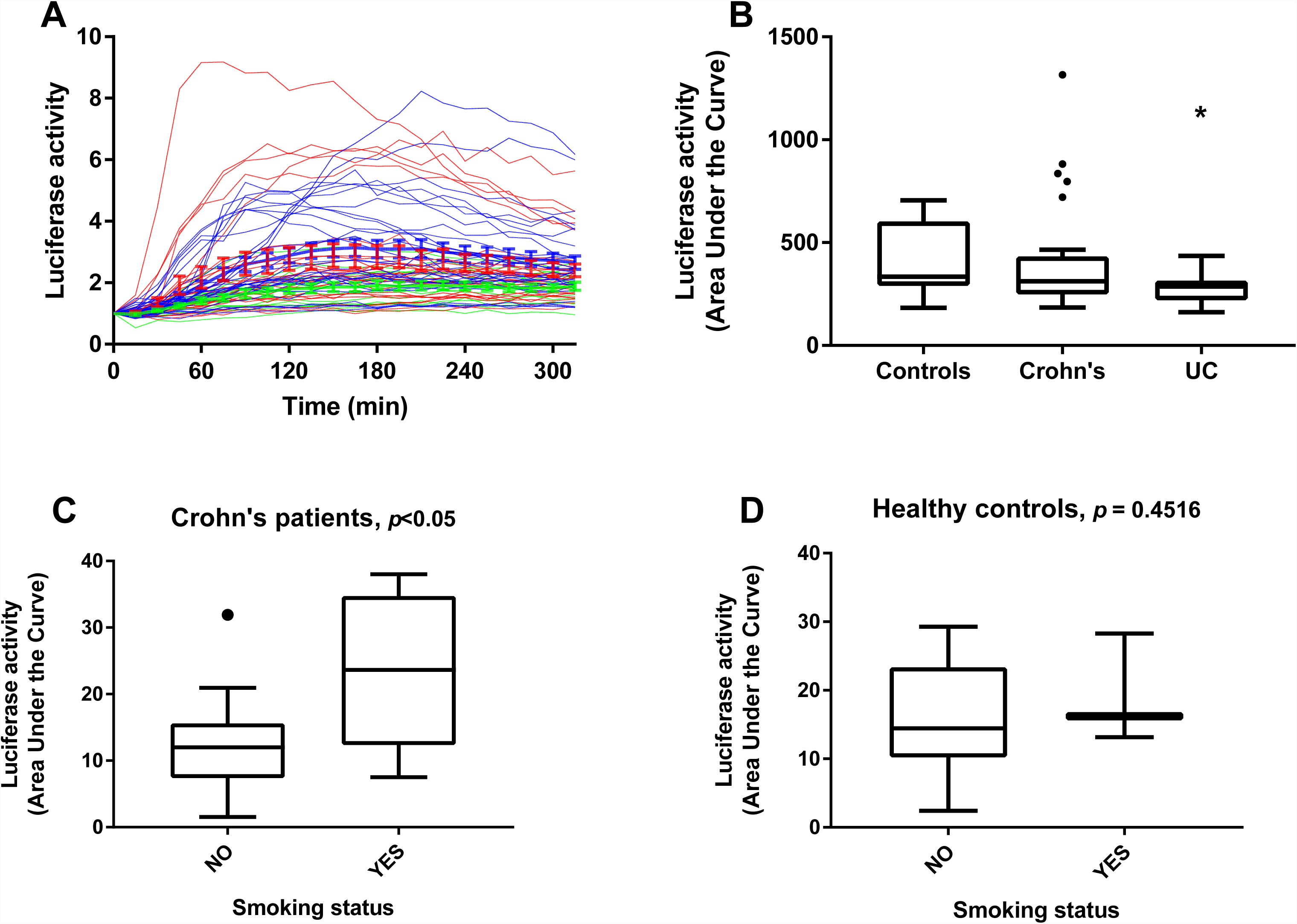
Luciferase activity in control and IBD patients. Luciferase activity from all patients screened represented as (A) dynamic, colour-coded graph (Blue=control, Red=CD and Green=UC) and (B) area under the curve (AUC) for control, CD and UC groups. NF-κB-regulated luciferase activity in Crohn’s patients (C) and control donors (D) based on smoking status. Statistical comparisons between disease types were made using the Kruskal-Wallis test, * denotes p<0.05. Statistical differences between smoking status were tested using the Mann-Whitney-U test, individual p-values are reported on each chart.

### Smoking status is the only independent factor which correlated with luciferase activity

In this relatively small cohort, our analysis demonstrated, after adjustment for other factors, that non-smokers had substantially lower luciferase activity than current smokers (coefficient: 7.68, 95% Confidence interval: 1.53; 13.83 and *p*=0.015) (Table 2). Patients with UC also had a trend towards lower luciferase activity than control patients, but this just failed to reach statistical significance (*p*=0.070).There was no significant difference between recruitment centres (*p*=0.771), and that neither gender (*p*=0.678), age (*p*=0.585), nor concomitant immunomodulatory (*p*=0.978), and/or biologic drug use (*p*=0.972), influenced the outcome of the luciferase activity assay.

**Table 2:**
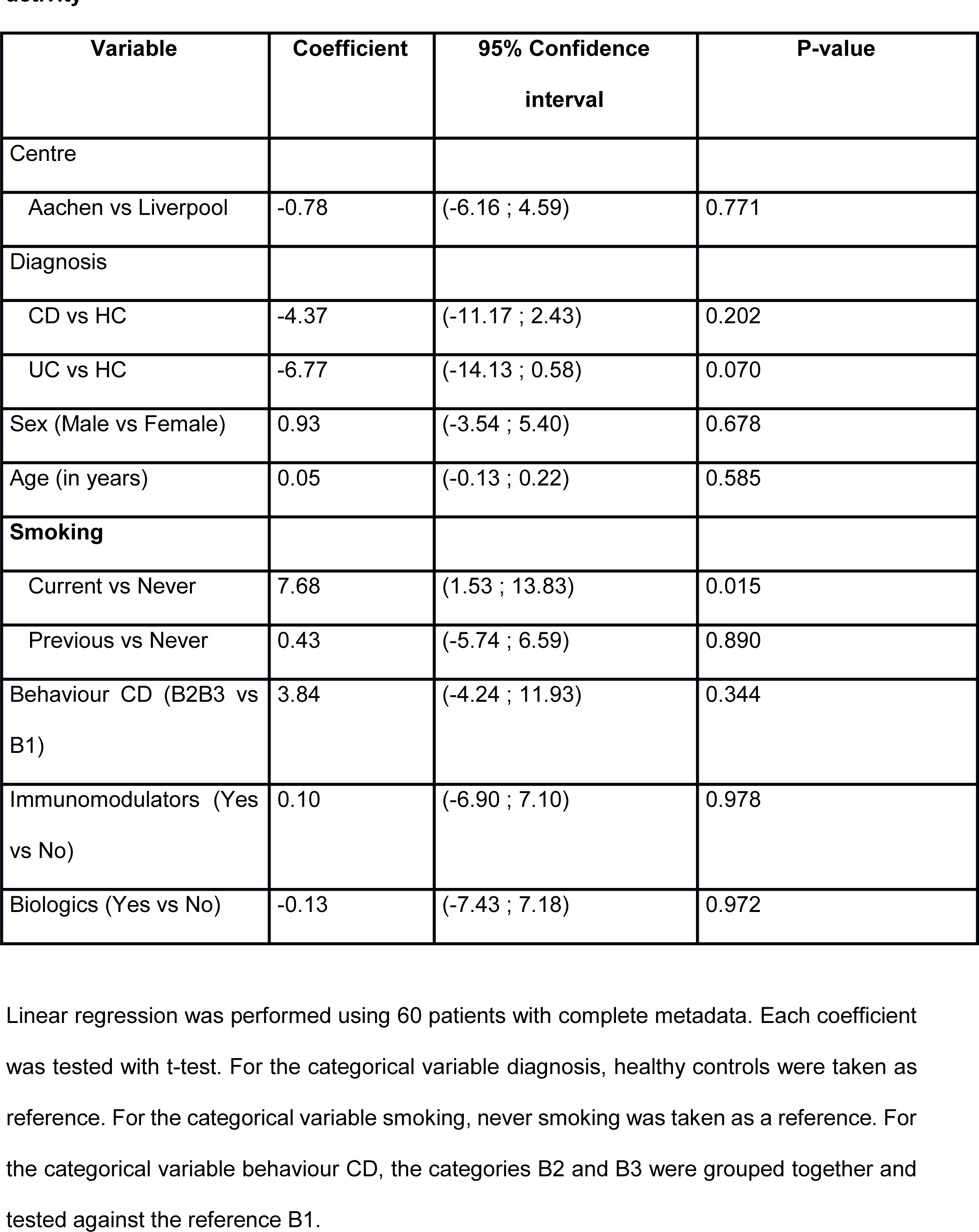
Demographic characteristics and associations with the AUC of the luciferase activity.

| Variable | Coefficient | 95% Confidence interval | P-value |
| --- | --- | --- | --- |
| Centre |  |  |  |
| Aachen vs Liverpool | -0.78 | (-6.16 ; 4.59) | 0.771 |
| Diagnosis |  |  |  |
| CD vs HC | -4.37 | (-11.17 ; 2.43) | 0.202 |
| UC vs HC | -6.77 | (-14.13 ; 0.58) | 0.070 |
| Sex (Male vs Female) | 0.93 | (-3.54 ; 5.40) | 0.678 |
| Age (in years) | 0.05 | (-0.13 ; 0.22) | 0.585 |
| **Smoking** |  |  |  |
| Current vs Never | 7.68 | (1.53 ; 13.83) | 0.015 |
| Previous vs Never | 0.43 | (-5.74 ; 6.59) | 0.890 |
| Behaviour CD (B2B3 vs B1) | 3.84 | (-4.24 ; 11.93) | 0.344 |
| Immunomodulators (Yes vs No) | 0.10 | (-6.90 ; 7.10) | 0.978 |
| Biologics (Yes vs No) | -0.13 | (-7.43 ; 7.18) | 0.972 |

Linear regression was performed using 60 patients with complete metadata. Each coefficient was tested with t-test. For the categorical variable diagnosis, healthy controls were taken as reference. For the categorical variable smoking, never smoking was taken as a reference. For the categorical variable behaviour CD, the categories B2 and B3 were grouped together and tested against the reference B1.

## Discussion

Inflammatory bowel disease has complex pathogenesis involving genetic susceptibility, intestinal microbiota, the host immune system and environmental factors such as diet, stress, smoking and hygiene (26). Defects in signalling pathways can lead to dysregulation of the inflammatory response that are crucial in the pathogenesis of IBD. One of the most studied pathways is the NF-κB signalling pathway which was first linked to IBD in 1998 (21). Since then, many laboratories have shown hyper-activation of the NF-κB signalling pathways in intestinal epithelial or immune cells from IBD patients (11,21,27-29).

Confocal microscopy has been used extensively in cell lines and primary cells for quantitative measurement of NF-κB nuclear translocation (8,25,30,31). In this study, we aimed to visualise NF-κB/p65 activation dynamics and this is the first report that has showed a correlation between NF-κB activation and IBD in primary human PBMDMs by confocal imaging. The p65 subunit showed cytoplasmic localization in resting cells and upon stimulation it exhibited rapid nuclear translocation. Notably, only a single p65 translocation event (rather than repeated oscillation) was observed within the 3h data acquisition period of confocal imaging. This is similar to previous analyses of murine bone marrow-derived macrophages from transgenic mice in response to either Lipid A or LPS (9,10). Interestingly, a significant difference was observed in the percentage of responding cells in samples from CD patients compared to controls. Although the reason for this differential response is unknown, it could explain why higher levels of activity were seen in cells from CD patients at a population level compared to healthy controls. There were no differences seen between donor groups with respect to the peak activation time, nor the intensity of the peak in response to LPS stimulation. There was however, marked difference in observed peak width in UC patient stimulated macrophages compared to those from healthy controls, (p<0.001) and to CD patients (p<0.05). The peak width defines the kinetics of NF-κB dwell time in the nucleus. We have shown previously that inhibition of nuclear export affects the dynamics of p65 localization as it is detected by confocal imaging (32). There is also a strong evidence that this defines cell specific patterns of gene expression (4,33) and it could correlate to disease status. Therefore, a smaller peak width implies that NF-κB likely stays for a shorter time in the nucleus and this could have an impact on gene transcription.

Taking these results further, we designed a screening strategy of human PBMDMs based on an NF-κB-regulated luciferase reporter *in vitro* assay. Luciferase reporter assays are widely used because they are convenient, relatively inexpensive, and give quantitative measurements instantaneously (34). Frozen, human PBMCs were used as starting material, and we optimized the culture conditions, the lentivirus infection conditions, and the luciferase assay protocol in order to achieve reproducible results. We then screened a novel cohort of subjects consisting of 65 donors from Liverpool and Aachen. Control donors from both clinical sites showed similar luciferase profiles and could reliably be used for comparison with samples from individuals affected by IBD. The luciferase activity profile was analysed and used to cluster the samples into three groups, showing Cluster 1 as low active group and Cluster 3 as high active group. Many cytokines are regulated by the same signalling pathways and NF-κB is a major pro-inflammatory transcription factor in immune cells. We therefore compared the amount of cytokines induced by LPS stimulation of cells from individuals in cluster 1 and Cluster 3. Those data together showed that there was a positive correlation between cytokine levels and endogenous NF-κB activation in human PBMDMs.

Static measurements of cytokines in matched intestinal biopsies and serum did not however demonstrate any correlation with either the NF-κB-based clustering or stimulated *in vivo* cytokine production.

In contrast, we observed that individuals who were assigned to cluster 1 based on their luciferase assay predominantly consisted of patients with UC, and this was validated when luciferase data were analysed based on disease status. This finding is in agreement with the initial single-cell imaging data and indeed linked the duration of nuclear localization to gene transcription as reflected by luciferase activity.

This observation is however different from previous reports of high NF-κB activity in UC patients (11, 21). This discrepancy may be due to the cell types that have been tested as well as the procedures that were used to prepare and assay the samples. On the other hand, individuals with CD showed a broader spectrum of luciferase activity. They represented the majority of samples in cluster 3, but also several samples in cluster 2. Metadata analysis of our cohort of patients with CD revealed that samples from active smokers had statistically significant higher NF-κB activity compared to non-smokers within this disease subgroup. This difference was not observed in the control donors who smoked but did not have IBD. Moreover, multivariate analysis of the entire cohort revealed that the only independent factor that predicted differences in luciferase activity was smoking.

There is an established strong association between smoking and CD, perhaps best demonstrated in the recent TOPPIC trial which identified smoking as the only factor which predicted post-operative recurrence in CD (35). Our study raises the hypothesis that the association of smoking and CD disease may be due to an effect of this combination on NF-κB signalling dynamics.

Our study did not investigate the mechanisms that underlie links between smoking and NF-κB activation, but we have shown that differences in our assay are sustained even after prolonged storage of cells, suggesting that it is unlikely to be a short-term influence of specific components of cigarette smoke that influences the differences we observe.

Cigarette smoking is reported to have long term epigenetic effects, some of which are permanent, whilst others are reversible, on smoking cessation (36). This mechanism may be one way in which smoking can have a sustained influence of NF-κB activity *in vitro*.

Amongst the top 10 research priorities identified by a recent James Lind alliance priority setting partnership was “What are the optimal markers/combinations of markers (clinical, endoscopic, imaging, genetics, other biomarkers) for stratification of patients with regards to a) disease course, b) monitoring disease activity and c) treatment response?” (37). The luciferase assay described in this manuscript shows promise both as a potential predictor of a diagnosis of ulcerative colitis, and as a potential stratification tool for future therapeutic studies that propose to target NF-κB activity. Before these uses can be established however, there will need to be extensive assessment of the assay in larger cohorts and these studies will need to be appropriately powered to determine how effectively the assay functions. A personalised medicine approach for IBD is attractive. The screening assay described in this study appears to be able to segregate IBD patients into 3 clusters and therefore has the potential to be used for further *in vitro* drug testing, specifically for patients who do not respond to therapy.

An additional and more pragmatic future goal may also be to use this assay in combination with epigenetic studies to attempt to identify links between inflammatory bowel diseases, NF-κB activation and smoking status.

## Methods

### Human study design - Ethics approval

We performed a study using healthy donors and IBD patients from the UK and Germany. In Liverpool (UK), patients attending for colonoscopy for any clinical indication were recruited. At University RWTH Aachen (Germany), individuals with an established diagnosis of IBD were recruited. Patient and healthy control blood samples and intestinal tissue biopsy specimens were obtained following informed consent. Patient and control samples were obtained following informed consent and with study approval from NRES Committee North West-Liverpool East (R&D 4910; REC 15/NW/0045) and the Regional Human Ethics Committee, Aachen, Germany (EK 235/13).

### Lentiviral NF-κB transcriptional activity vectors

NF-κB transcriptional activity was monitored using a lentiviral construct (κB-NLSluc) that expresses firefly luciferase under the control of the classical NF-κB promoter (38). For imaging, the human p65 sequence was amplified from p65-dsRedXp (32) and C-terminally fused with AmCyan protein using a previously described lentiviral vector (9). Lentivirus production was carried out as per (39).

### Confocal imaging of p65-AmCyan lentivirus-transfected Human Peripheral Blood Mononuclear-Derived Macrophages (PBMDMs)

Fresh PBMCs were obtained from whole blood taken from patients recruited in Liverpool. PBMCs (4×10^6^ cells) were plated in 35mm μ-plate imaging dishes (Ibidi GmbH; Martinsried, Germany) and differentiated (Supplementary information: Methods). On day 4, macrophages were infected by addition of p65-AmCyan lentivirus into the culture medium. After 72h incubation, media was removed, fresh medium containing supplements (but no M-CSF) added and cells were imaged 24h later. Cells were stained for 1h with 10ng/mL Hoechst 33342 (Sigma), medium changed and cells rested for 1h prior to imaging. Cells were imaged using a Zeiss LSM880 confocal microscope system equipped with a cell incubation unit maintained at 37**°**C, in a humidified atmosphere of 5% CO_2_. p65-AmCyan nuclear fluorescence was detected (excitation λ 458nm, emission λ 489nm) and quantified using CellTracker software (40). Basal readings were obtained for 30min and cells then stimulated with 200ng/mL Lipid A (Sigma). Cells with a high starting variance in the absence of stimulation (standard deviation >10) were excluded from analyses. All measures performed 200min after stimulation were also excluded. Graphs were analysed based on the fact that a responsive cell is defined by a peak that is more than 2-fold above mean baseline values obtained before stimulation. The first peak width was defined as the length of time between the first point where nuclear fluorescence was ≥2-fold above baseline and the subsequent time point when nuclear fluorescence fell 2-fold below that of baseline.

### Human Peripheral Blood Mononuclear cell-Derived Macrophages (PBMDMs) – κB-NLSluc luciferase assay

Frozen peripheral blood mononuclear cell (PBMCs) isolated from peripheral venous blood were thawed and differentiated to PBMDMs (Supplementary Information: Methods). Transduced PBMDMs were cultured in 24-well plates (OptiPlate-24, White Opaque 24-well Microplate) in 0.4mL medium containing 1mM luciferin. Cells were stimulated with 200 ng/mL LPS and luminescence detected over time in a CO_2_ Lumistar Omega luminometer. Post-assay, pro-viral copies were measured by qPCR, using a Lenti-X™ Provirus Quantitation Kit (Clontech; Oxford, UK). For *in vitro* stimulation, other ligands used included human recombinant Interleukin-1 beta [IL-1β] (PeproTech), Flagellin FliC from *Salmonella* typhimurium (NovusBio; Littleton CO, USA), muramyl-dipeptide [MDP] (InvivoGen; Toulouse, France) and LPS extracted using modified phenol/water method (41) from IBD mucosa-associated *E. coli* isolates, LF82 and LF10 (42,43) (Supplementary Information: Methods).

### Human intestinal tissue specimens

Human biopsies (sigmoid colon and terminal ileum) obtained at colonoscopy following informed consent were stored at −80°C. After thaw on ice, they were lysed in 50μL sterile PBS by high-speed shaking (2 × 2min at 30Hz; TissueLyser II (QIAGEN)). After centrifugation at 10,000x*g* for 15min, each cleared tissue lysate was stored at −80°C.

### Cytokine measurements

PBMDMs were stimulated with 200ng/mL LPS for 20h and culture medium was harvested and stored at −80°C for cytokine quantification. Cytokines were measured using the V-PLEX Proinflammatory Panel 1 Human kit (Meso Scale Discovery; Rockville MD, USA). Total protein was measured in biopsy lysates using the bicinchoninic acid (BCA) assay (Pierce).

### Statistical approaches

see Supplementary Information: Methods

## Acknowledgment

The authors acknowledge the support of all SysMedIBD partners. The ileal CD mucosa-associated *E. coli* clinical isolates were a kind gift to BJC from the late Prof. Arlette Darfeuille-Michaud; INSERM Unit, Clermont-Ferrand, France. The human p65-AmCyan construct was provided by Dr James Bagnall, University of Manchester, UK.

## Contributors

SP, HE and MB: acquisition of data; SP, MB, FB and HE: analysis and interpretation of data; FB, BJC: data analysis, bioinformatics and statistical analysis; RH: human sample preparation and isolation of PBMCs; LS: lentiviral preparation; DGS: confocal imaging; BJC: purification of LPS from clinical *E. coli* isolates; CP, DMP, DJ, GS: provision of patient samples; SP, MB, BJC: drafted manuscript; MHRW, BJC, CP, DJ, GS, VMDS, PP, RH, WM, DMP: critical revision of the manuscript for important intellectual content; MHRW, BJC, DMP, VMDS, DJ, CP and WM: obtained funding. All the authors approved the final manuscript submission.

## Funding

This study was supported by the European SysMedIBD FP7 Programme (Agreement number 305564). The funder provided no input to the study design nor in the collection, analysis and interpretation of data.

## Competing interests

VDS is a director and shareholder of LifeGlimmer GmbH. FB has received salary from LifeGlimmer GmbH.

## References

1. Stevens C, Walz G, Singaram C, Lipman ML, Zanker B, Muggia A, et al. Tumor necrosis factor-alpha, interleukin-1 beta, and interleukin-6 expression in inflammatory bowel disease. Dig Dis Sci. 1992;37(6):818–26.

2. Strober W, Fuss IJ. Proinflammatory cytokines in the pathogenesis of inflammatory bowel diseases. Gastroenterology. 2011;140(6):1756–67.

3. Hayden MS, Ghosh S. NF-kappaB, the first quarter-century: remarkable progress and outstanding questions. Genes & development. 2012;26(3):203–34.

4. Ashall L, Horton CA, Nelson DE, Paszek P, Harper CV, Sillitoe K, et al. Pulsatile stimulation determines timing and specificity of NF-kappaB-dependent transcription. Science. 2009;324(5924):242–6.

5. Sung M-H, Li N, Lao Q, Gottschalk RA, Hager GL, Fraser IDC. Switching of the Relative Dominance Between Feedback Mechanisms in Lipopolysaccharide-Induced NF-κB Signaling. 2014;7(308):ra6–ra.

6. Tay S, Hughey JJ, Lee TK, Lipniacki T, Quake SR, Covert MW. Single-cell NF-kappaB dynamics reveal digital activation and analogue information processing. Nature. 2010;466(7303):267–71.

7. Kellogg RA, Tay S. Noise Facilitates Transcriptional Control under Dynamic Inputs. Cell. 2015;160(3):381–92.

8. Adamson A, Boddington C, Downton P, Rowe W, Bagnall J, Lam C, et al. Signal transduction controls heterogeneous NF-[kappa]B dynamics and target gene expression through cytokine-specific refractory states. Nat Commun. 2016;7.

9. Bagnall J, Boddington C, England H, Brignall R, Downton P, Alsoufi Z, et al. Quantitative analysis of competitive cytokine signaling predicts tissue thresholds for the propagation of macrophage activation. Sci Signal. 2018;11(540).

10. Cheng Z, Taylor B, Ourthiague DR, Hoffmann A. Distinct single-cell signaling characteristics are conferred by the MyD88 and TRIF pathways during TLR4 activation. Science Signaling. 2015;8(385).

11. Atreya I, Atreya R, Neurath MF. NF-kappaB in inflammatory bowel disease. J Intern Med. 2008;263(6):591–6.

12. Neurath MF, Fuss I, Schurmann G, Pettersson S, Arnold K, Muller-Lobeck H, et al. Cytokine gene transcription by NF-kappa B family members in patients with inflammatory bowel disease. Ann N Y Acad Sci. 1998;859:149–59.

13. Rogler G, Brand K, Vogl D, Page S, Hofmeister R, Andus T, et al. Nuclear factor kappaB is activated in macrophages and epithelial cells of inflamed intestinal mucosa. Gastroenterology. 1998;115(2):357–69.

14. Gelbmann CM, Leeb SN, Vogl D, Maendel M, Herfarth H, Scholmerich J, et al. Inducible CD40 expression mediates NFkappaB activation and cytokine secretion in human colonic fibroblasts. Gut. 2003;52(10):1448–56.

15. Merga YJ, O’Hara A, Burkitt MD, Duckworth CA, Probert CS, Campbell BJ, et al. Importance of the alternative NF-kappaB activation pathway in inflammation-associated gastrointestinal carcinogenesis. Am J Physiol Gastrointest Liver Physiol. 2016;310(11):G1081–90.

16. Burkitt MD, Hanedi AF, Duckworth CA, Williams JM, Tang JM, O’Reilly LA, et al. NF-kappaB1, NF-kappaB2 and c-Rel differentially regulate susceptibility to colitis-associated adenoma development in C57BL/6 mice. J Pathol. 2015;236(3):326–36.

17. Martinez-Montiel MP, Casis-Herce B, Gomez-Gomez GJ, Masedo-Gonzalez A, Yela-San Bernardino C, Piedracoba C, et al. Pharmacologic therapy for inflammatory bowel disease refractory to steroids. Clin Exp Gastroenterol. 2015;8:257–69.

18. Peyrin-Biroulet L, Lemann M. Review article: remission rates achievable by current therapies for inflammatory bowel disease. Aliment Pharmacol Ther. 2011;33(8):870–9.

19. Roda G, Jharap B, Neeraj N, Colombel JF. Loss of Response to Anti-TNFs: Definition, Epidemiology, and Management. Clin Transl Gastroenterol. 2016;7:e135.

20. Atreya R, Neurath MF. Mechanisms of molecular resistance and predictors of response to biological therapy in inflammatory bowel disease. Lancet Gastroenterol Hepatol. 2018;3(11):790–802.

21. Schreiber S, Nikolaus S, Hampe J. Activation of nuclear factor kappa B inflammatory bowel disease. Gut. 1998;42(4):477–84.

22. Murray PJ, Allen JE, Biswas SK, Fisher EA, Gilroy DW, Goerdt S, et al. Macrophage Activation and Polarization: Nomenclature and Experimental Guidelines. Immunity. 2014;41(1):14–20.

23. Raetz CR, Reynolds CM, Trent MS, Bishop RE. Lipid A modification systems in gram-negative bacteria. Annual review of biochemistry. 2007;76:295–329.

24. Turner DA, Paszek P, Woodcock DJ, Nelson DE, Horton CA, Wang Y, et al. Physiological levels of TNFalpha stimulation induce stochastic dynamics of NF-kappaB responses in single living cells. J Cell Sci. 2010;123(Pt 16):2834–43.

25. Kardynska M, Paszek A, Smieja J, Spiller D, Widlak W, White MRH, et al. Quantitative analysis reveals crosstalk mechanisms of heat shock-induced attenuation of NF-kappa B signaling at the single cell level. Plos Comput Biol. 2018;14(4).

26. Dabritz J, Menheniott TR. Linking immunity, epigenetics, and cancer in inflammatory bowel disease. Inflamm Bowel Dis. 2014;20(9):1638–54.

27. Smythies LE, Shen R, Bimczok D, Novak L, Clements RH, Eckhoff DE, et al. Inflammation anergy in human intestinal macrophages is due to Smad-induced IkappaBalpha expression and NF-kappaB inactivation. J Biol Chem. 2010;285(25):19593–604.

28. Naiki Y, Michelsen KS, Zhang W, Chen S, Doherty TM, Arditi M. Transforming growth factor-beta differentially inhibits MyD88-dependent, but not TRAM- and TRIF-dependent, lipopolysaccharide-induced TLR4 signaling. J Biol Chem. 2005;280(7):5491–5.

29. Pasparakis M. IKK/NF-kappaB signaling in intestinal epithelial cells controls immune homeostasis in the gut. Mucosal Immunol. 2008;1 Suppl 1:S54–7.

30. Noursadeghi M, Tsang J, Haustein T, Miller RF, Chain BM, Katz DR. Quantitative imaging assay for NF-kappaB nuclear translocation in primary human macrophages. J Immunol Methods. 2008;329(1-2):194–200.

31. Sillitoe K, Horton C, Spiller DG, White MR. Single-cell time-lapse imaging of the dynamic control of NF-kappaB signalling. Biochem Soc Trans. 2007;35(Pt 2):263–6.

32. Nelson DE, Ihekwaba AE, Elliott M, Johnson JR, Gibney CA, Foreman BE, et al. Oscillations in NF-kappaB signaling control the dynamics of gene expression. Science. 2004;306(5696):704–8.

33. Nelson G, Paraoan L, Spiller DG, Wilde GJ, Browne MA, Djali PK, et al. Multi-parameter analysis of the kinetics of NF-kappaB signalling and transcription in single living cells. J Cell Sci. 2002;115(Pt 6):1137–48.

34. Fan F, Wood KV. Bioluminescent assays for high-throughput screening. Assay Drug Dev Technol. 2007;5(1):127–36.

35. Mowat C, Arnott I, Cahill A, Smith M, Ahmad T, Subramanian S, et al. Mercaptopurine versus placebo to prevent recurrence of Crohn’s disease after surgical resection (TOPPIC): a multicentre, double-blind, randomised controlled trial. Lancet Gastroenterol Hepatol. 2016;1(4):273–82.

36. Joehanes R, Just AC, Marioni RE, Pilling LC, Reynolds LM, Mandaviya PR, et al. Epigenetic Signatures of Cigarette Smoking. Circ Cardiovasc Genet. 2016;9(5):436–47.

37. Ma C, Jairath V, Khanna R, Feagan BG. Investigational drugs in phase I and phase II clinical trials targeting interleukin 23 (IL23) for the treatment of Crohn’s disease. Expert Opin Investig Drugs. 2018:1–12.

38. Brignall R, Cauchy P, Bevington SL, Gorman B, Pisco AO, Bagnall J, et al. Integration of Kinase and Calcium Signaling at the Level of Chromatin Underlies Inducible Gene Activation in T Cells. The Journal of Immunology. 2017;199(8):16.

39. Bagnall J, Boddington C, Boyd J, Brignall R, Rowe W, Jones NA, et al. Quantitative dynamic imaging of immune cell signalling using lentiviral gene transfer. Integr Biol-Uk. 2015;7(6):713– 25.

40. Shen H, Nelson G, Nelson DE, Kennedy S, Spiller DG, Griffiths T, et al. Automated tracking of gene expression in individual cells and cell compartments. J R Soc Interface. 2006;3(11):787– 94.

41. Apicella MA, Griffiss JM, Schneider H. Isolation and characterization of lipopolysaccharides, lipooligosaccharides, and lipid A. Methods Enzymol. 1994;235:242–52.

42. Boudeau J, Glasser AL, Masseret E, Joly B, Darfeuille-Michaud A. Invasive ability of an Escherichia coli strain isolated from the ileal mucosa of a patient with Crohn’s disease. Infect Immun. 1999;67(9):4499–509.

43. Masseret E, Boudeau J, Colombel JF, Neut C, Desreumaux P, Joly B, et al. Genetically related Escherichia coli strains associated with Crohn’s disease. Gut. 2001;48(3):320–5.

