## Supplementary information for "Macrophage-specific NF-κB activation dynamics can segregate inflammatory bowel disease patients"

### Materials and Methods

#### Extraction of LPS from IBD-mucosa associated *Escherichia coli*

The protocol used for the isolation of LPS from *E. coli* was based on the modified phenol/ water technique as per (26). Two ileal IBD mucosa-associated *E. coli* clinical isolates, LF82 and LF10 (27, 28), were grown overnight in Luria-Bertani broth. Bacteria were centrifuged and washed in sterile endotoxin free water and lyophilised overnight. Lyophilised *E. coli* (5g) were ground to a fine powder and re-suspended in 25mL of 50mM sodium phosphate-5mM EDTA pH 7 and allowed to hydrate completely. The suspension was stirred in a shearing mixer at top speed for 1min before 100mg lysozyme (Sigma) was added and the suspension was then incubated overnight, continuously stirring, at 4$^{o}$C. The following day, the suspension was incubated at 37$^{o}$C for 20min and stirred in a shearing mixer for 3min. The suspension volume was increased to 100mL by adding 50mM sodium phosphate-20mM $MgCl_2$ pH 7 containing 1μg/mL ribonuclease A and deoxyribonuclease I (Sigma), followed by incubation for 1h at 37$^{o}$C. The suspension was placed in a 70$^{o}$C water bath until temperatures equilibrated and an equal volume of preheated (70$^{o}$C) 90%w/v phenol was added and thoroughly mixed. The resulting mixture was rapidly cooled by stirring for 15min in an ice-water bath. The phenol-bacterial suspension was centrifuged at 4$^{o}$C at 18,000x*g* for 15min – a sharp interface occurs between the aqueous and phenol layers. The aqueous layer containing LPS was removed by aspiration and dialysed at 4$^{o}$C against frequent changes of endotoxin-free water. The LPS was lyophilised and stored at -80$^{o}$C.

**Human Peripheral Blood Mononuclear cell-Derived Macrophages (PBMDMs) – Isolation and *in vitro* differentiation**

Peripheral venous blood (10mL) was immediately heparinized (unfractionated heparin sodium, at 5U/mL; Wockhardt UK Ltd; Wrexham; Wales). Each sample was mixed 1:2 with sterile phosphate-buffered saline pH7.3 (PBS), layered over 20mL Ficoll-Paque™ plus (Thermo-Fisher Scientific; Paisley, UK) and centrifuged at 400x*g* for 40min at room temperature. Peripheral blood mononuclear cells (PBMCs) were aspirated, washed with sterile PBS, resuspended in 1mL freezing medium (88%v/v FCS (Sigma) + 12%v/v DMSO (Sigma)) and stored at -80ºC. Frozen isolated Peripheral Blood Mononuclear cells (PBMCs) were thawed and plated ($4x10^6$ cells/well) in 6-well plates (Nunclon Vita surface; Thermo Fisher Scientific) in 3mL differentiation medium [RPMI-1640, 10%v/v FCS, 10mM HEPES (Sigma), 1mM sodium pyruvate (Thermo Fisher Scientific), 1X MEM non-essential amino acids (Thermo Fisher Scientific), 10U/mL penicillin/ 10mg/mL streptomycin/ 2mM L-glutamine (Sigma) and 50ng/mL human macrophage colony-stimulating factor (M-CSF, Peprotech; London, UK)]. On day 1, adherent cells were washed and 3mL differentiation medium was added for a further 3-6d depending on the experiment. Following differentiation, 70-80% of the adherent macrophages (PBMDMs) expressed characteristic macrophage cell-surface markers (CD11b+, CD14 low). PBMDM cultures were infected with κB-NLSluc lentivirus on day 4 and were incubated for 24h. The volume of the lentivirus used was optimized per virus batch to achieve the highest level of transduction without causing cell death.

**Statistical analysis**

Mann-Whitney-Wilcoxon tests and Pearson's chi-squared proportional test were performed to compare patient demographics. T-tests and correlational tests were performed to compare the luciferase activation to clinical parameters. P-values were adjusted for multiple testing of outcomes using the false discovery rate method. Multivariate analyses were performed using the R language, and univariate analyses were performed using either the R language or GraphPad Prism v7.0. For cluster analysis, K-medoids algorithm was applied to stratify the patients with respect to their $\log_2$ fold-change of luciferase activity. The number of clusters was optimised with the average silhouette width criterion. In order to understand whether the association between disease status and luciferase activity was real, or confounded by other clinical or demographic factors, we performed a linear regression analysis of key variables against luciferase activity. Mann-Whitney U test was used for univariate analysis of cytokine levels in LPS-stimulated PBMDMs, in serum and in biopsy lysates. Fisher's exact test was used to analyse the responding cells obtained by the confocal imaging data. Kruskal-Wallis test was used to analyse the peak width in the responding cells obtained by the confocal imaging data.

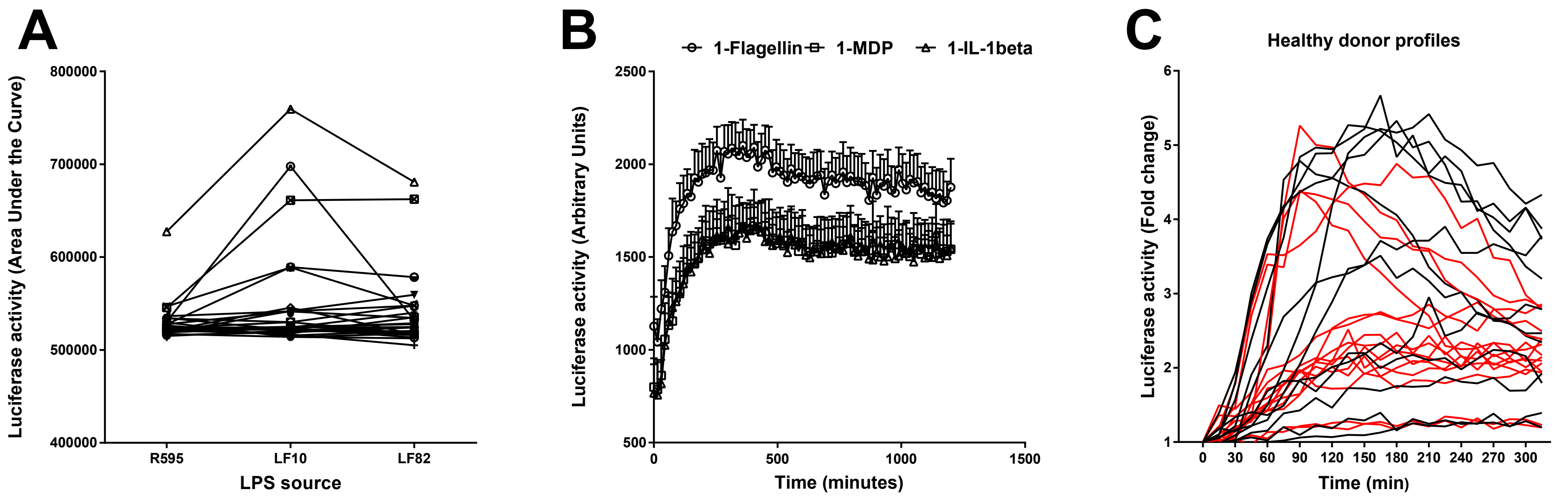

A
800000
700000
600000
500000
400000
Luciferase activity (Area Under the Curve)
R595
LF10
LF82
LPS source
B
1-Flagellin
1-MDP
1-IL-1beta
2500
2000
1500
1000
500
Luciferase activity (Arbitrary Units)
0
500
1000
1500
Time (minutes)
C
Healthy donor profiles
6
5
4
3
2
1
Luciferase activity (Fold change)
0
30
60
90
120
150
180
210
240
270
300
Time (min)

**Figure S1**: Luciferase activity of PBMDMs stimulated by various ligands. PBMDMs show similar NF-κB luciferase activity, represented as area under the curve, as a response to 200ng/mL LPS from various sources (A). PBMDMs were stimulated with 100ng/mL Flagellin (-o-), 500ng/mL MDP (-□-) and 20ng/mL IL-1β (-Δ-) and luciferase activity was measured over time, as shown in (B). Luciferase activity profiles in LPS-stimulated PBDMs from healthy donors from Liverpool (black) and Aachen (red) (C).

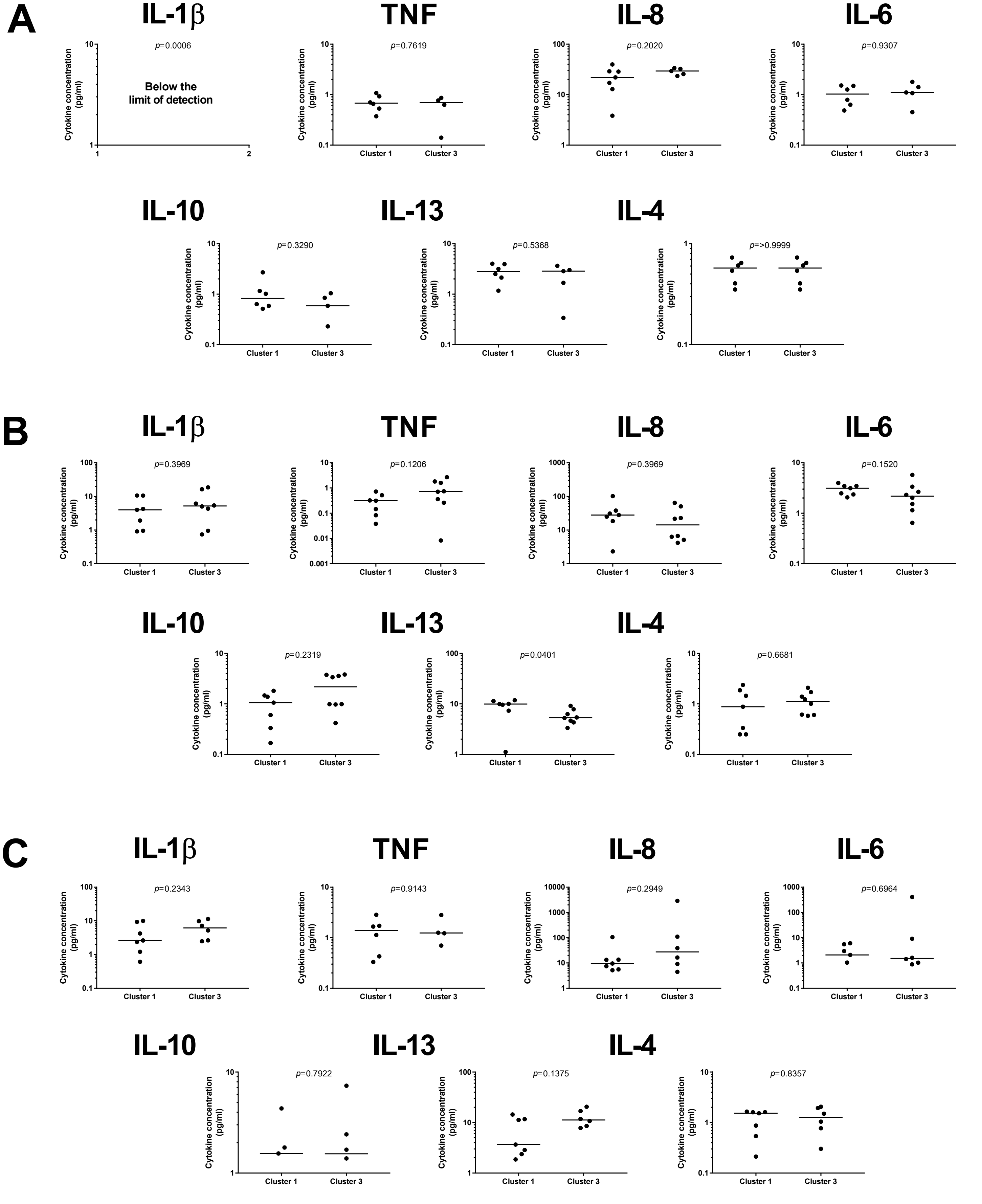
A
IL-1β
p=0.0006
Below the limit of detection
TNF
p=0.7619
IL-8
p=0.2020
IL-6
p=0.9307
IL-10
p=0.3290
IL-13
p=0.5368
IL-4
p=>0.9999
B
IL-1β
p=0.3969
TNF
p=0.1206
IL-8
p=0.3969
IL-6
p=0.1520
IL-10
p=0.2319
IL-13
p=0.0401
IL-4
p=0.6681
C
IL-1β
p=0.2343
TNF
p=0.9143
IL-8
p=0.2949
IL-6
p=0.6964
IL-10
p=0.7922
IL-13
p=0.1375
IL-4
p=0.8357
Cytokine concentration (pg/ml)
Cluster 1
Cluster 3

**Figure S2**: No correlation between dynamic and static measurements in cytokine levels. Comparison of cytokine levels produced by LPS-stimulated PBMDMs (dynamic) versus cytokine levels detected in serum (top panel), sigmoid colon biopsy lysates (middle panel) and terminal ileum biopsy lysates (bottom panel). Results of sum-of-squares linear regression analysis (r2 and F-test p- values) are reported on each graph.

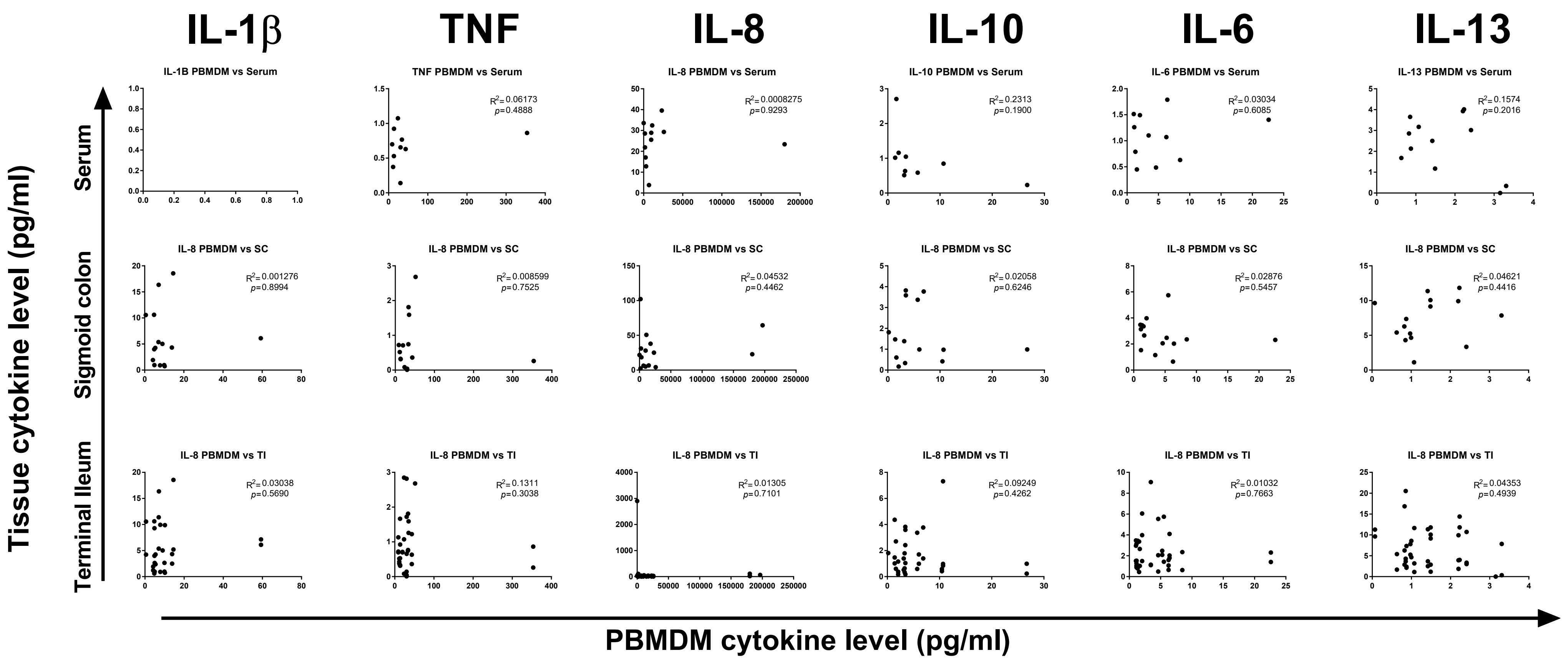

IL-1β
TNF
IL-8
IL-10
IL-6
IL-13
Tissue cytokine level (pg/ml)
Serum
Sigmoid colon
Terminal Ileum
IL-1B PBMDM vs Serum
TNF PBMDM vs Serum
R2=0.06173
p=0.4888
IL-8 PBMDM vs Serum
R2=0.0008275
p=0.9293
IL-10 PBMDM vs Serum
R2=0.2313
p=0.1900
IL-6 PBMDM vs Serum
R2=0.03034
p=0.6085
IL-13 PBMDM vs Serum
R2=0.1574
p=0.2016
IL-8 PBMDM vs SC
R2=0.001276
p=0.8994
IL-8 PBMDM vs SC
R2=0.008599
p=0.7525
IL-8 PBMDM vs SC
R2=0.04532
p=0.4462
IL-8 PBMDM vs SC
R2=0.02058
p=0.6246
IL-8 PBMDM vs SC
R2=0.02876
p=0.5457
IL-8 PBMDM vs SC
R2=0.04621
p=0.4416
IL-8 PBMDM vs TI
R2=0.03038
p=0.5690
IL-8 PBMDM vs TI
R2=0.1311
p=0.3038
IL-8 PBMDM vs TI
R2=0.01305
p=0.7101
IL-8 PBMDM vs TI
R2=0.09249
p=0.4262
IL-8 PBMDM vs TI
R2=0.01032
p=0.7663
IL-8 PBMDM vs TI
R2=0.04353
p=0.4939
PBMDM cytokine level (pg/ml)

**Figure S3**: No correlation between dynamic and static measurements in cytokine levels. Comparison of cytokine levels produced by LPS-stimulated PBMDMs (dynamic) versus cytokine levels detected in serum (top panel), sigmoid colon biopsy lysates (middle panel) and terminal ileum biopsy lysates (bottom panel). Results of sum-of-squares linear regression analysis (r2 and F-test p-values) are reported on each graph.

**Macrophage-specific NF-κB activation dynamics can segregate inflammatory bowel disease patients.** Stamatia Papoutsopoulou, Michael D. Burkitt, François Bergey, Hazel England, Rachael Hough, Lorraine Schmidt, David G Spiller, Mike HR White, Pawel Paszek, Dean A. Jackson, Gernot Sellge, D. Mark Pritchard, Vitor A.P. Martins Dos Santos, Barry J. Campbell, Werner Müller, Chris S. Probert.

**Table S1**: **Percentage of responding cells in samples from each patient as assessed by confocal imaging of PBMDM cultures infected with lentivirus vector expressing the human p65-AmCyan protein and stimulated *in vitro* with 200ng/mL Lipid A.**

| **Healthy controls**<br><br>**N=6** | **CD**<br><br>**N=5** | **UC**<br><br>**N=3** |
|---|---|---|
| 25.0 | 16.7 | 60.0 |
| 70.4 | 33.3 | 20.0 |
| 30.0 | 85.7 | 90.0 |
| 30.0 | 63.6 | |
| 10.0 | 66.7 | |
| 57.1 | | |

The total percentage of responding cells was calculated based on the assumption that a responsive cell is defined by a peak that is more than 2-fold above baseline values (i.e. mean of the values before stimulation). CD = Crohn’s disease, UC = ulcerative colitis
